# Genetic Influence underlying Brain Connectivity Phenotype: A Study on Two Age-Specific Cohorts

**DOI:** 10.1101/2021.08.23.457353

**Authors:** Shan Cong, Xiaohui Yao, Linhui Xie, Jingwen Yan, Li Shen, for the Alzheimer’s Disease Neuroimaging Initiative

## Abstract

**Background:** Human brain structural connectivity is an important imaging quantitative trait for brain development and aging. Mapping the network connectivity to the phenotypic variation provides fundamental insights in understanding the relationship between detailed brain topological architecture, function, and dysfunction. However, the underlying neurobiological mechanism from gene to brain connectome, and to phenotypic outcomes, and whether this mechanism changes over time, remain unclear.

**Methods:** This study analyzes diffusion weighted imaging data from two age-specific neuroimaging cohorts, extracts structural connectome topological network measures, performs genome-wide association studies (GWAS) of the measures, and examines the causality of genetic influences on phenotypic outcomes mediated via connectivity measures.

**Results:** Our empirical study has yielded several significant findings: 1) It identified genetic makeup underlying structural connectivity changes in the human brain connectome for both age groups. Specifically, it revealed a novel association between the minor allele (G) of rs7937515 and the decreased network segregation measures of the left middle temporal gyrus across young and elderly adults, indicating a consistent genetic effect on brain connectivity across the lifespan. 2) It revealed rs7937515 as a genetic marker for body mass index (BMI) in young adults but not in elderly adults. 3) It discovered brain network segregation alterations as a potential neuroimaging biomarker for obesity. 4) It demonstrated the hemispheric asymmetry of structural network organization in genetic association analyses and outcome-relevant studies.

**Discussion:** These imaging genetic findings underlying brain connectome warrant further investigation for exploring their potential influences on brain-related diseases, given the significant involvement of altered connectivity in neurological, psychiatric and physical disorders.

**Impact Statement:** The genetic architecture underlying brain connectivity, and whether this mechanism changes over time, remain largely unknown. To understand the inter-individual variability at different life stages, this study performed genome-wide association studies of brain network connectivity measures from two age-specific neuroimaging cohorts, and identified a common association between the minor allele (G) of rs7937515 and decreased network segregation measures of the left middle temporal gyrus. The mediation analysis further elucidated neurobiological pathway of brain connectivity mediators linking the genes *FAM86C1*/*FOLR3* with body mass index. This study provided new insights into the genetic mechanism of inter-regional connectivity alteration in the brain.

## 1 Introduction

Brain structural connectivity is a major organizing principle of the nervous system. Estimating interregional neural connectivity, reconstructing geometric structure of fiber pathways, and mapping the network connectivity to corresponding inter-individual variabilities provide fundamental insights in understanding detailed brain topological architecture, function and dysfunction. A large body of research has been devoted to extracting and investigating macro-scale brain networks from diffusion-weighted imaging (DWI) data (Bertolero et al., 2019; Elsheikh et al., 2020; van den Heuvel et al., 2019; Jiang et al., 2019; Xie et al., 2018), and various behavioural, neurological and neuropsychiatric disorders have been linked to the disrupted brain connectivity (van den Heuvel et al., 2019; Jiang et al., 2019). As structural changes of brain connectivity are phenotypically associated with massive complex traits across different categories, the brain-wide connectome has been extensively studied.

It is worth noting that human brain connectome re-configures its network structure dynamically and adaptively in response to genetic, lifestyle, environmental factors (Cauda et al., 2018; Cohen and D’Esposito, 2016), brain development and aging (Alloza et al., 2018; Sala-Llonch et al., 2015; Varangis et al., 2019). However, the underlying neurobiological mechanism from gene to brain connectome, and to cognitive and behavioral outcomes, and whether this mechanism changes over time, remain unclear. To bridge this gap, we perform a genetic study of brain connectome phenotypes on two different age-specific cohorts: one contains healthy young adults (age: 28.7 *±* 3.6), and the other contains elderly participants (age: 73.8 *±* 7.0). Our goal is to identify genetic factors affecting brain connectivity and examine their consistency and discrepancy between these two age-specific groups.

Emerging advances in multimodal brain imaging, high throughput genotyping and sequencing techniques provide exciting new opportunities to ultimately improve our understanding of brain structure and neural dynamics, their genetic architecture and their influences on cognition and behavior (Shen and Thompson, 2020). Present studies investigating direct associations among human connectomics, genomics and clinical phenotyping are primarily focused on four aspects: (1) estimating genetic heritability of basic connectome measures such as number of fibers, length of fibers and fractional anisotropy (FA) (Elliott et al., 2018; Jahanshad et al., 2013; Thompson et al., 2013); (2) discovering pairwise univariate associations between single nucleotide polymorphisms (SNPs) and imaging phenotypic traits such as above mentioned basic connectome measures at each edge (Jahanshad et al., 2013; Karwowski et al., 2019) and white matter properties at each voxel (Alloza et al., 2018; Guo et al., 2020; Kochunov et al., 2010); (3) discovering pairwise univariate associations between SNPs and clinical phenotypes such as cognitive or behavioral outcomes (Elsheikh et al., 2020; Jahanshad et al., 2013); and (4) discovering pairwise univariate associations between basic connectome measures and clinical phenotypes (van den Heuvel et al., 2019; Jiang et al., 2019).

Among the studies mentioned above, there exist two major limits. First, these studies were conducted based on basic connectome measures such as number of fibers, length of fibers and FA, but the complex-network attributes were overlooked, which included network segregation, integration, centrality and resilience and important network components such as hubs, communities, and rich clubs (Sporns, 2013). These attributes were extensively adopted to detect network integration and segregation, quantitatively measure the centrality of network regions and pathways, characterize patterns of local anatomical circuitry, and test resilience of networks to insult (Rubi-nov and Sporns, 2010). Second, these studies performed analyses by examining the association between an independent variable (e.g. SNP) and a dependent variable (e.g. cognitive or behavioral outcome), without taking into consideration the mediator(s) linking these variables (Baron and Kenny, 1986). Mediation analysis can help identify the underlying mechanism of outcome-relevant genetic effects implicitly mediated by neuroimaging phenotypes (e.g. connectome measures). Of note, mediation analysis requires the independent variable to be significantly associated with both the dependent variable and the mediator. This makes applying it in brain neuroimaging studies a challenge due to the modest effect size of an individual genetic variant on both behavioral and imaging phenotypes (Cong et al., 2018; Saykin et al., 2015), as well as limited size of the sample with all diagnostic, imaging and genetic data available.

With the demand of measuring complex-network attributes, a few recent genome-wide association studies (GWAS) (Bertolero et al., 2019; Elsheikh et al., 2020) recognized the first problem mentioned above and adopted quantitative measurement approaches for complex-network attributes , and treated the attributes as neuroimaging traits for the explorations of complex imaging genomic associations. They successfully identified a number of loci susceptible for Alzheimer’s Disease (Elsheikh et al., 2020), and demonstrated the associations between loci and segregated network patterns, which may be involved in brain development, evolution, and disease (Bertolero et al., 2019). However, a notable limitation is that these studies only focus on the brain networks of either young or elderly participants, as a result, their study outcomes are lack of validations in multiple data sets. Since there is an age-related discrepancy for genetic effects on human connectome alterations across lifespan (Varangis et al., 2019), it remains an under-explored topic to examine genetic consistency and discrepancy for complex-network attributes among cohorts different in age. Another factor that may cause discrepancy in the network architecture is the hemispheric asymmetry (Jiang et al., 2019), and the hemispheric asymmetry of network organization has been linked to development processes (Zhong et al., 2017) and neuropsychiatric disorders (Sun et al., 2017). It remains a challenge to understand the genetic basis for the network attributes of two hemispheres as they may be distinctively correlated to cognition level, physical and psychological development.

Among a large amount of complex-network attributes, it has been well documented in recent literatures (Cohen and D’Esposito, 2016; Xie et al., 2018) that segregation of neural information such as modularity, transitivity, clustering coefficients and local efficiency represent the connectivity of local network communities that are intrinsically densely connected and strongly coupled. A converging evidence (Cohen and D’Esposito, 2016; Karwowski et al., 2019) is shown that local, within-network communication is critical for motor execution, whereas integrative, between-network communication is critical for measuring connectome (Bertolero et al., 2019). Thus, network segregation is thought to be essential for describing and understanding of complex neural connectome systems (Sporns, 2013). In addition, segregation measures are highly reliable and heritable network attributes (Xie et al., 2018), and these measures has been linked to the disruption of neural network connectivity in brain development, evolution, disease (Bertolero et al., 2019; Cohen and D’Esposito, 2016; Mak et al., 2016), and immunodeficiency (Bell et al., 2018). Given the importance of network segregation, in this study, we first focus on quantifying measures of network segregation, analyzing heritability of segregation measures and performing genetic association analyses by treating them as neuroimaging traits. Then, our next priority is to explore the genetic basis for the rest of the complex-network attributes (e.g. integration, centrality and resilience).

To overcome the challenges mentioned above, this study aims to develop and implement computational and statistical strategies for a systematic characterization of structural connectome optimized for imaging genetic studies, and to determine genetic basis of structural connectome. Specifically, the framework is organized and described in Figure 1, and the primary goals are to address the following six critical issues: (a) construction of basic network connectivity with diffusion tractography, (b) systematic extraction of complex-network attributes, (c) heritability analysis of complex-network attributes, (d) genome-wide association studies of quantitative endopheno-types, (e) examination of mediation effect that intermediately bridges genes and outcomes, and (f) identification of outcome-relevant neuroimaging biomarkers. Given the enormously broad scope of brain connectome, our focus is on studying (1) static tractography-based structural connectome and complex-network attributes characterizing segregation, integration, centrality and resilience; (2) genetic consistency and discrepancy for complex-network attributes among cohorts different in age; and (3) mediation effects of network attributes on outcome-relevant genetics.

**Figure 1:**
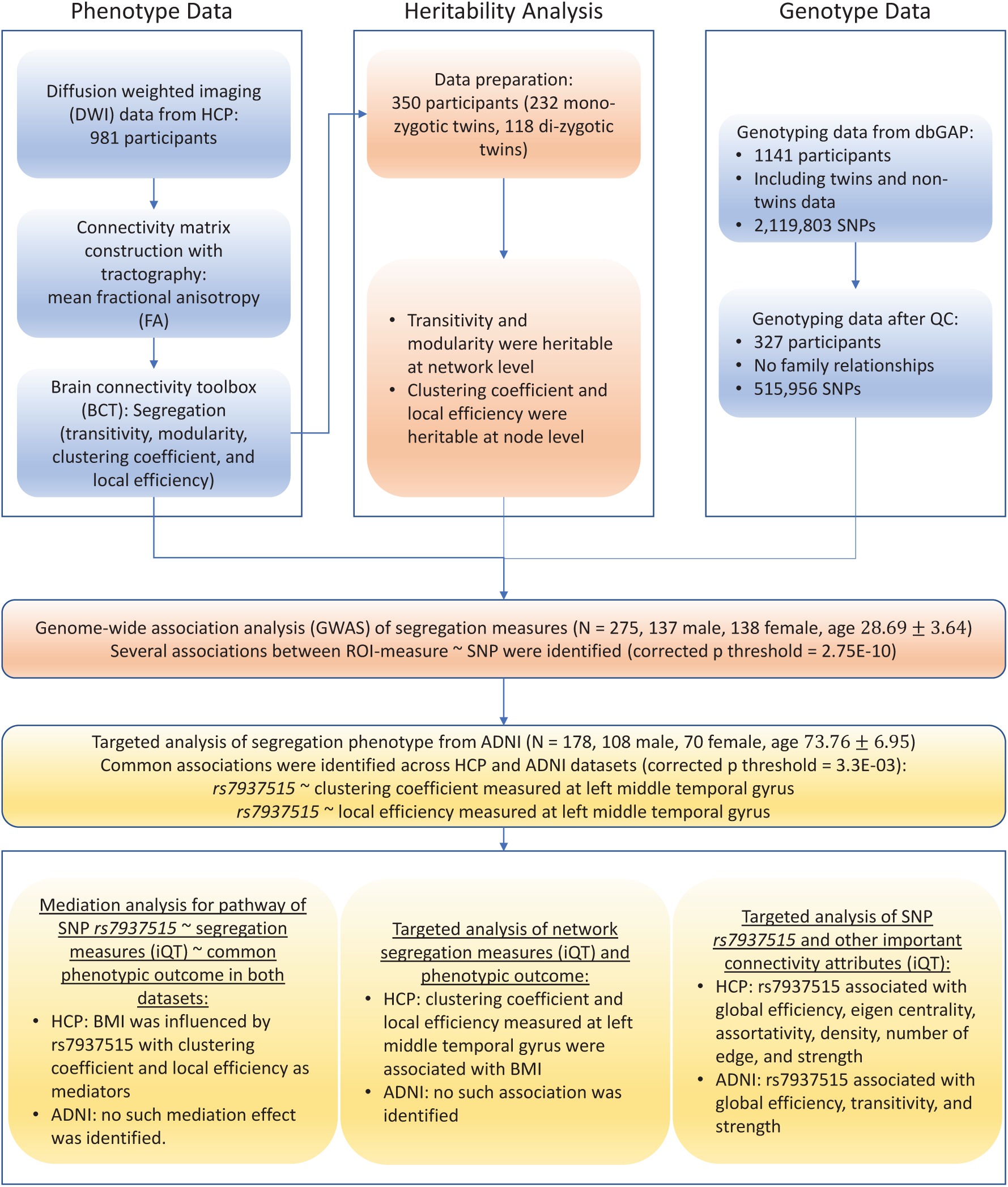
Flowchart of brain connectome GWAS design. Abbreviations: SNPs, single nucleotide polymorphisms; ADNI, Alzheimer’s Disease Neuroimaging Initiative; HCP, Human Connectome Project; dbGaP, database of Genotypes and Phenotypes; QC, quality control; ROI, region of interest; iQT: imaging quantitative trait; BMI, body mass index.

The major contributions of this study are fivefold:

- New challenges in human connectome: we elucidate the neurobiological pathway from SNPs to brain connectome, and to phenotypic outcomes. By integrating connectomics and genetics, this study provides new genetic mechanism insights into understanding detailed brain topological architecture, and encoding (or mapping) inter-regional connectivity in the genome.
- New genetic insights for brain phenotype: we validate the study outcomes by examining genetic consistency and discrepancy for complex-network attributes between young adult cohort and elderly adult cohort, which illustrates genetic basis for human connectome in different life stages.
- Biological findings: we treat network segregation measures as imaging quantitative traits (iQT), and demonstrate that body mass index (BMI, as a phenotypic outcome) is influenced by a locus rs7937515 with network segregation attributes (e.g. clustering coefficient and local efficiency) measured at the left middle temporal gyrus as mediators, which reveals the intermediate effects of brain connectivity in the pathway of outcome-relevant genetics.
- Biological findings: we discover network segregation as important neuroimaging biomaker for BMI and weight-related issues, and illustrate the importance of the left middle temporal gyrus for BMI.
- Biological findings: we demonstrate the hemispheric asymmetry of structural network organization in genetic association analyses and outcome-relevant studies.

## 2 Materials and Methods

### 2.1 Study datasets

With the purpose of examining genetic consistency and discrepancy for complex-network attributes between young and elderly adults, and illustrating genetic basis for human connectome in different life stages, our analysis was respectively conducted on Human Connectome Project (HCP) database for young adults and Alzheimer’s Disease Neuroimaging Initiative (ADNI) database for elderly adults.

#### 2.1.1 HCP young adult dataset

HCP (Van Essen et al., 2013) is a major endeavor to map macroscopic human brain circuits and their relationship to behavior in a large population. It charts the neural pathways that underlie brain function and behavior, by acquiring and analyzing human brain connectivity from high-quality neuroimaging data in healthy young adults. The HCP datasets serve as a key resource for the neuroscience research community, as it provides valuable resources for characterizing human brain connectivity and function, their relationship to behavior, and their heritability and genetic underpinnings, which enables discoveries of how the brain is wired and how it functions in different individuals.

#### 2.1.2 ADNI elderly adults data set

Alzheimer’s Disease Neuroimaging Initiative (ADNI) database was initially launched in 2004 as a public-private partnership, and led by the Principal Investigator Michael W. Weiner, MD. One primary aim of ADNI has been to examine whether serial imaging biomarkers extracted from MRI, positron emission tomography (PET), other biological markers, and clinical and neuropsychological assessment can be combined to measure the progression of mild cognitive impairment (MCI) and early AD. For up-to-date information, see www.adni-info.org.

### 2.2 Demographics

We initially downloaded 981 subjects from HCP database, including a part of twin subjects, then one individual from each family was randomly selected and excluded. As a result, 275 unrelated participants were selected for further population-based genetic analyses. ADNI data were collected by selecting the participants who had both genotype data and baseline DWI data at their first visit, family relationship was also removed in the same way as described above for HCP data filtration. Detailed characteristic information and the number of subjects in each data cohort are shown in Table 1. In this study, we analyzed a total of 275 participants (age: 28.7 *±* 3.6; gender: 137 male, 138 female; education: 15.1 *±* 1.6) from the HCP database, and a total of 178 participants (age: 73.8 *±* 7.0; gender: 108 male, 70 female; education: 16.0 *±* 2.8) from the ADNI database. This study was approved by institutional review boards of all participating institutions, and written informed consent was obtained from all participants or authorized representatives.

**Table 1.**
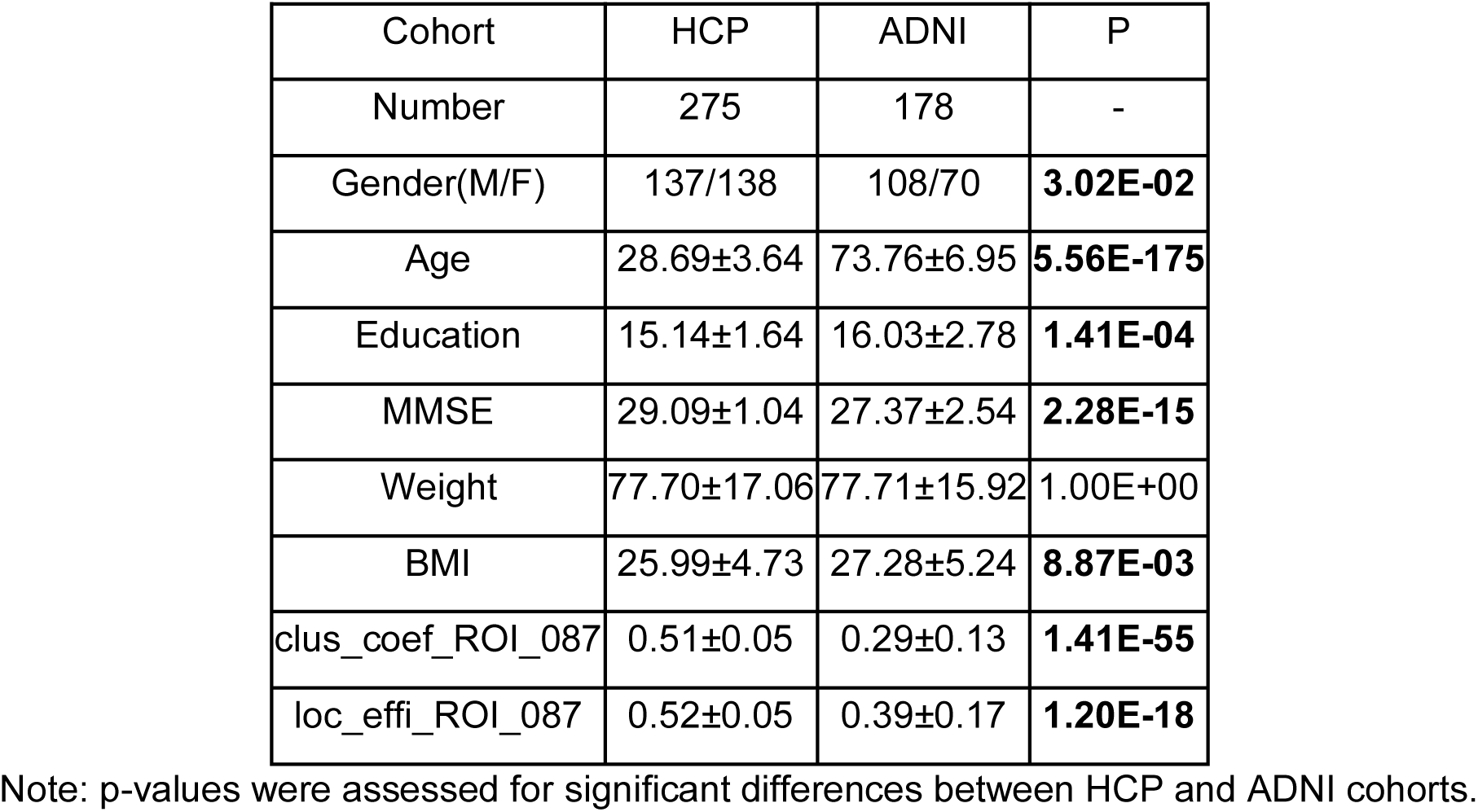
Participant characteristics in genetic association of brain connectomic network.

### 2.3 Genotyping data acquisition and processing

#### 2.3.1 HCP young adults dataset

HCP samples were genotyped using MEGA array with PsychChip and ImmunoChip content. 1,141 genotype data was downloaded from dbGAP. Individuals were retained if they had a genotype rate *≥* 0.98. Samples with evidence of relatedness (IBD estimate *≥* 0.25) or heterozygosity rate *±*3 standard deviations from the mean were also excluded. SNPs were excluded if they had a minor allele frequency (MAF) *≤* 0.05, call rate *≤* 98% or a Hardy-Weinberg *P ≤* 1.0*×*10^−6^, and SNPs with no “rs” number were further excluded. Following quality control process, the number of genotyping data reduced to 327, we then checked the missing data by matching subjects information between phenotype and genotype data. As a result, this study comprised a total of 327 unrelated subjects and 515,956 SNPs.

#### 2.3.2 ADNI elderly adults dataset

Genotyping data were quality-controlled, imputed and combined as described in (Cong et al., 2020; Yao et al., 2020, 2019). Briefly, genotyping was performed on all ADNI participants following the manufacturer’s protocol using blood genomic DNA samples and Illumina GWAS arrays (610-Quad, OmniExpress, or HumanOmni2.5-4v1) (Saykin et al., 2010). Quality control was performed in PLINK v1.90 (Purcell et al., 2007) using the following criteria: 1) call rate per marker *≥* 95%, 2) minor allele frequency (MAF) *≥* 5%, 3) Hardy Weinberg Equilibrium (HWE) test P *≤* 1.0E-6, and 4) call rate per participant *≥* 95%. In total, 5,574,300 SNPs were included for further targeted genetic association analysis.

### 2.4 Tractography and network construction

#### 2.4.1 Tractography

We downloaded high spatial resolution DWI data and genotype data from both HCP and ADNI databases. DWI data from HCP was processed following the MRtrix3 guidelines (Tournier et al., 2012), detailed procedures have been previously reported (Xie et al., 2018) and are briefly described below: (1) generating a tissue-segmented image; (2) estimating the multi-shell multi-tissue response function and performing the multi-shell multi-tissue constrained spherical deconvolution; (3) generating the initial tractogram and applying the successor of Spherical-deconvolution Informed Filtering of Tractograms (SIFT2) methodology (Smith et al., 2015); and (4) mapping the SIFT2 output streamlines onto the MarsBaR automated anatomical labeling (AAL) atlas (Tzourio-Mazoyer et al., 2002) with 90 ROIs to produce the structural connectome with edge value equal to the mean fractional anisotropy (FA).

DWI data from ADNI was acquired following the scanning protocols described in (Elsheikh et al., 2020), and processed following the procedures discussed in (Yan et al., 2018). Tractography was performed in Camino (Cook et al., 2006) based on white matter fiber orientation distribution function (ODF). As Camino adopted a deterministic approach, streamlines were modeled with a multi-tensor modeling approach (voxels fitted up to three fiber orientations, this way accounting for most of the fiber-crossings) of the ODF data. To generate a comparable tractography, the streamlines were also mapped onto AAL atlas with 90 ROIs to produce the structural connectome with edge value equal to the mean FA.

#### 2.4.2 Network construction

Network was created and defined by connectivity matrix *M* where *M_ij_* stores the connectivity measure between ROIs *i* and *j*. As described in the previous section, we considered FA for defining *M_ij_* . Since the diffusion tensor is a symmetric 3 *×* 3 matrix, it can be described by its eigenvalues (*λ*_1_, *λ*_2_ and *λ*_3_) and eigenvectors (*v*_1_, *v*_2_ and *v*_3_) for tractography analysis. FA at edge-level is an index for the amount of diffusion asymmetry within a voxel, defined in terms of its eigenvalues:

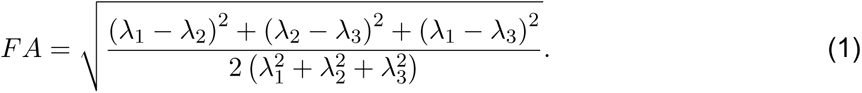

Thus, mean FA is a normalized measure of the fraction of the tensor’s magnitude due to anisotropic diffusion, corresponding to the degree of anisotropic diffusion or directionality.

### 2.5 Complex-network attributes

With an undirected and weighted connectivity matrix *M* (defined in Section 2.4.2), we assessed a comprehensive set of network features such as segregation (e.g. transitivity, clustering coefficients, local efficiency and modularity), integration (e.g. global efficiency), centrality (e.g. eigen centrality) and resilience (e.g. assortativity) of the structural connectome using Brain Connectivity Toolbox (BCT) (Rubinov and Sporns, 2010). Given the importance and priority of segregation measures in this study, we only introduced the definitions of segregation measures, and the definitions of the rest complex-network attributes were explained in (Rubinov and Sporns, 2010).

For the following sub-sections, we define *N* as the set of all nodes in the network, *n* as the number of nodes, *t_i_* as geometric mean of triangles around node 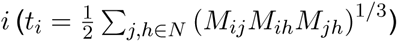, *k_i_* as weighted degree of *i* (*k_i_* = Σ*_j∈N_ M_ij_*), *a_ij_* as the connection status between *i* and *j* (*a_ij_* = 1 when link (*i, j*) exists, *a_ij_* = 0 otherwise), *d_ij_* as shortest weighted path length between *i* and *j* 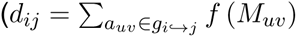, where *f* is a map from weight to length and *g_i→j_* is the shortest weighted path between *i* and *j*).

#### 2.5.1 Transitivity

Transitivity measures the ratio of triangles to triplets in the network. By following the definition in (Newman, 2003):

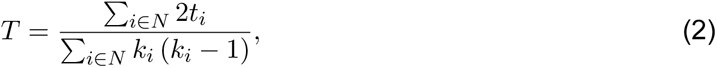

where *T* is the transitivity measured at network level.

#### 2.5.2 Clustering coefficient

Clustering coefficient measures the degree to which nodes in a network tend to cluster together by evaluating the fraction of triangles around a node and is equivalent to the fraction of node’s neighbors that are neighbors of each other. By following the definition in (Onnela et al., 2005):

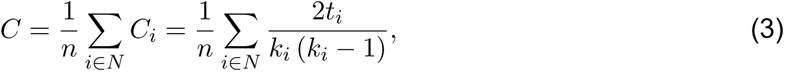

where *C_i_* is the clustering coefficient of node *i* and *C* is the clustering coefficient measured at network level.

#### 2.5.3 Local efficiency

Local efficiency measures the efficiency of information transfer limited to neighboring nodes by evaluating the global efficiency computed on node neighborhoods. By following the definition in (Latora and Marchiori, 2001):

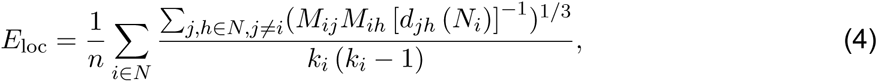

where *E*_loc_ is the local efficiency of node *i*, and *d_jh_* (*N_i_*) is the length of the shortest path between *j* and *h*, that contains only neighbors of *i*.

#### 2.5.4 Modularity

Modularity measures network segregation into distinct networks, and it is a statistic that quantifies the degree to which the network may be subdivided into such clearly delineated groups (Newman, 2006):

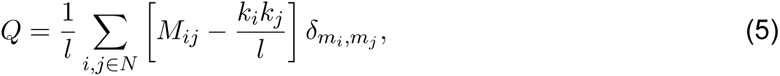

where *Q* is the modularity measured at network level, *m_i_* is the module containing node *i*, and *δ_mi,mj_* = 1 if *m_i_* = *m_j_*, and 0 otherwise.

### 2.6 Heritability Analysis

Heritability analysis focused on identifying highly heritable measures of structural brain networks, and it was a commonly adopted and critical measurement to describe properties of the inheritance of iQT. An iQT such as network attributes must be heritable, which was defined as the proportion of phenotypic variance due to genetic differences between individuals (Jørstad and Næevdal, 1996). In this study, we estimated heritability of four segregation measures from twin subjects in the HCP young adult cohort (*N* = 350, 232 mono-zygotic twins, 118 di-zygotic twins) and SOLAR-Eclipse software (Kochunov et al., 2015) was employed for this task. The inputs to this software included phenotype traits, covariates measures and a kinship matrix indicating the pairwise relationship between twin individuals. A maximum likelihood variance decomposition method was applied to estimate the variance explained by additive genetic factors and environmental factors respectively. The output from SOLAR-Eclipse included heritability (h2), standard error and the corresponding significance p-value for each feature. We estimated the heritability of connectomic features, including transitivity, clustering coefficients, local efficiency and modularity. Since many previous studies had reported the effect of age (linear nonlinear), gender and their interactions on structural brain connectivity (Burzynska et al., 2010; Gong et al., 2011; Lopez-Larson et al., 2011; Zhao et al., 2015), all heritability analyses were performed with age, age^2^, sex, age*×*sex and age^2^*×*sex as covariates. In addition, we extracted the total variance explained by all covariate variables.

### 2.7 Brain connectome genetic association analysis

GWAS on the brain network segregation measures of the 90 ROIs were performed using linear regression under an additive genetic model in PLINK v1.90 (Purcell et al., 2007). Age, gender and education were included as covariates. Our GWAS was focused on analyzing the following network segregation measures: (1) modularity and transitivity, which were network-level measures; and (2) clustering coefficient and local efficiency, which were node-level measures. As a result, in total, we have 2+90*×*2 = 182 measures. Our post-hoc analysis used Bonferroni correction for correcting the genome-wide significance (GWS) for the number of quantitative traits (i.e., 5E-8/182 = 2.75E-10).

Genetic findings of the segregation measures from HCP young adult dataset were treated as genotypic candidates and segregation measures at specific ROIs as phenotypic candidates, we further examined in ADNI elderly adult dataset regarding their associations. By validating the genetic findings from HCP data using ADNI participants, we examined genetic consistency and discrepancy for network segregation attributes between young and elderly adults, which illustrated the consistency and discrepancy of genetic basis for human connectome in different life stages.

In addition, the validated genetic findings were used to further explore connectivity variances with all important complex-network attributes excepting segregation measures such as integration (e.g. global efficiency and network density), centrality (e.g. eigen centrality) and resilience (e.g. assortativity), and we examined the targeted genetic basis on certain brain ROIs (e.g. middle temporal gyrus). As previously stated, linear regression models were used. In particular, we applied additive genetic models implemented in PLINK v1.90, with age, gender, education as covariates.

### 2.8 Mediation analysis

To examine the causal assumption, we followed (Baron and Kenny, 1986) to perform standard mediation analysis to identify the mediated effect, and we treated iQTs (e.g. network segregation measures) as mediating variables, which intermediately linked the pathological path from genetic findings to clinical phenotypes. Specifically, we constructed a set of candidate SNPs which were found significantly associated to segregation measures in both young and elderly participants, and we constructed a set of candidate clinical phenotyping information by extracting overlapped clinical outcomes collected in both HCP and ADNI databases. We then employed the mediation model to detect the indirect effect of loci on clinical outcomes via iQT.

Let *y* be the dependent variable which was a clinical outcome in our study, *x* be the independent variable which was a candidate SNP, *z* be the covariates (age, gender and education), and *M* be the mediating variable which was brain iQT. Mediation analysis were performed by following the three steps listed below.

**Step 1:** We adopted linear regression to regress the clinical outcome *y* against SNP *x* while controlling for *z*:

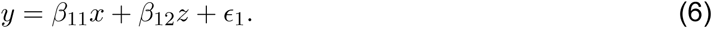

In the first step, the mediation assumption is satisfied if the coefficient *β*_11_ is statistically significant after correction for multiple comparisons. In this study, we employed Bonferroni correction, which lead the significant p-value threshold to: 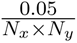, where *N_x_* was the total number of candidate SNPs, and *N_y_* was the total number of clinical outcomes.

**Step 2:** We adopted linear regression to regress the mediating variable *M* against SNP *x* while controlling for *z*:

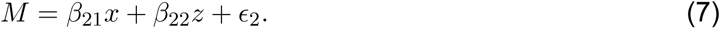

In the second step, the mediation assumption is satisfied if the coefficient *β*_21_ is statistically significant after Bonferroni correction for multiple comparisons, which lead the significant p-value threshold to: 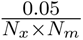, where *N_m_* is the total number of iQTs.

**Step 3:** We adopted linear regression to regress clinic outcome *y* against SNP *x* and the mediating variable *M* while controlling for *z*:

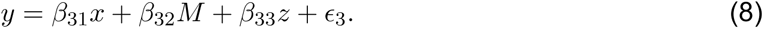

The third step was performed only when the mediating variable successfully passed the second step. We again employed Bonferroni correction for multiple comparisons, which lead the significant p-value threshold to: 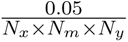. In the third step, the mediation assumption is satisfied if the coefficient *β*_32_ is statistically significant and *β*_32_ *< β*_11_ at the same time, which demonstrates an indirect mediated effect was identified between outcome *y* and SNP *x* mediated through iQT *M* .

### 2.9 Outcome-relavent brain connectome association analysis

To discover the outcome-relevant biomarkers which mapped brain connectivity alterations to cognitive or behavioral outcomes, we performed pairwise univariate association analysis between network segregation attributes and outcome data. We selected BMI and Mini-Mental State Examination (MMSE) as outcomes as they were the only measures available in both HCP and ADNI. We used linear regression to regress the phenotypic outcomes against network segregation measures for both HCP and ADNI datasets, and explored outcome-relevant brain neuroimaging biomarkers. By comparing young and elderly participants, we illustrated the consistency and discrepancy of human brain connectome in different ages regarding on BMI and MMSE variations.

## 3 Results

### 3.1 Heritability of network segregation

As illustrated in Figure 1, we examined segregation measures estimated at both network-level and node-level prior to GWAS. All of the segregation measures such as clustering coefficients (node-level), local efficiencies (node-level), transitivity (network-level) and modularity (network-level) showed significantly high heritability after Bonferroni correction (p*<* 0.05/182 = 2.75E-04). The mean (*±*std) heritability of 182 segregation measures (h2 score) was 0.81(*±*0.05), and more detailed results of heritability analysis were listed in Supplementary Table. We included all 182 segregation measures in the subsequent GWAS analysis.

### 3.2 GWAS of network segregation in HCP young adults

In the HCP cohort, genome-wide associations between 515, 956 SNPs and 182 structural network segregation measures were assessed under the additive genetic model and controlled for age, gender and education. GWAS identified 20 significant associations between 10 SNPs and 7 segregation measures (Table 2), after correcting the genome-wide significance (GWS) for the number of phenotypes using Bonferroni method (i.e., p*<* 5E-08/182 = 2.75E-10). Respectively shown in Figure 2 were Manhattan plots of GWAS results of clustering coefficient and local efficiency measured in left middle temporal gyrus. GWAS of HCP data showed high consistency for clustering coefficient and local efficiency, where nine SNP-ROI associations were discovered for these two segregation measures. After Bonferroni correction, there was no significant finding for the network level segregation measures (i.e., transitivity and modularity).

**Figure 2:**
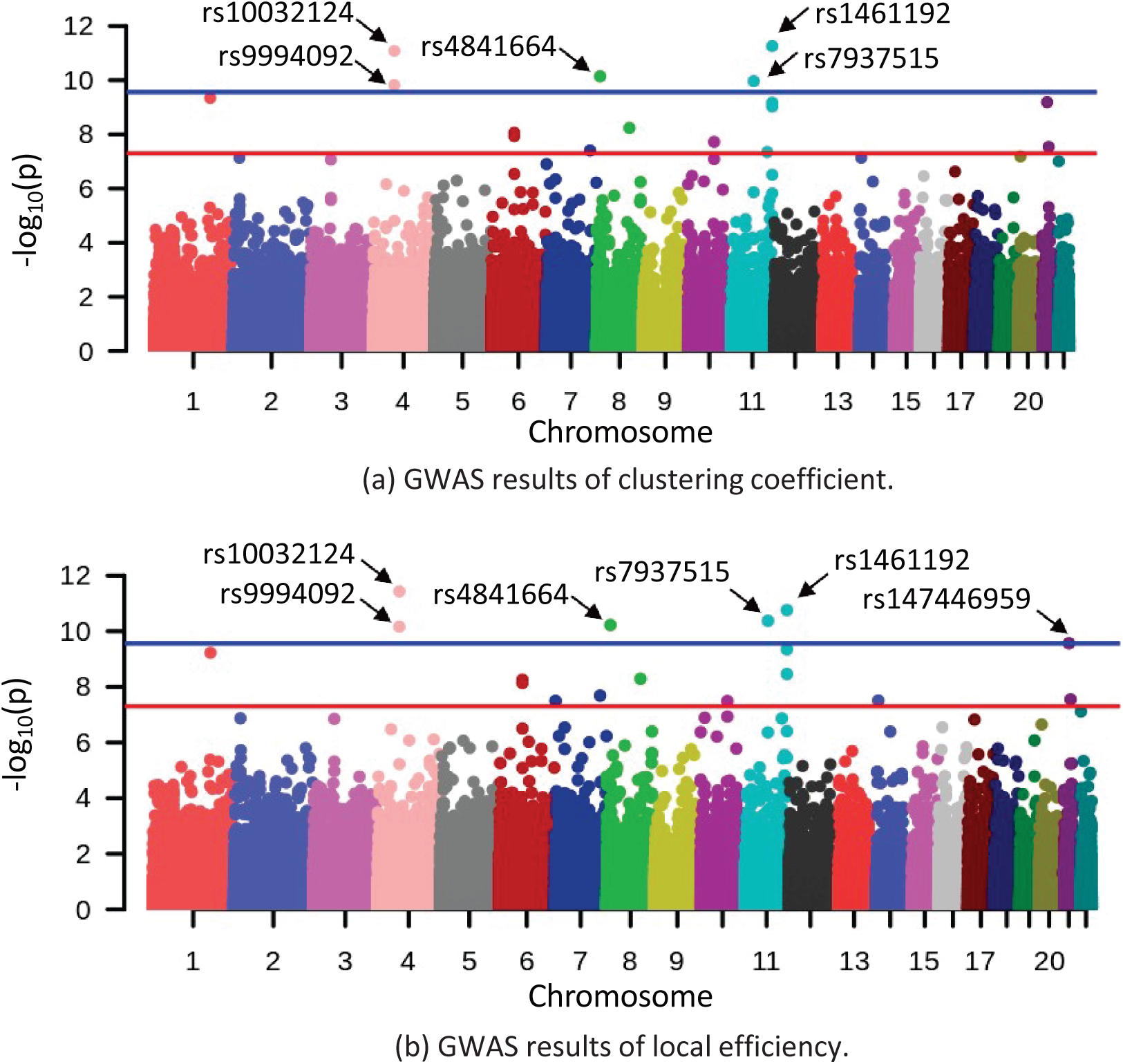
Manhattan plot of GWAS results in the HCP dataset. Top and bottom plots show the GWAS results of clustering coefficient and local efficiency on left middle temporal gyrus, respectively. Red and blue lines correspond to the p-value of 5E-08 and 2.75E-10, respectively.

**Table 2.**
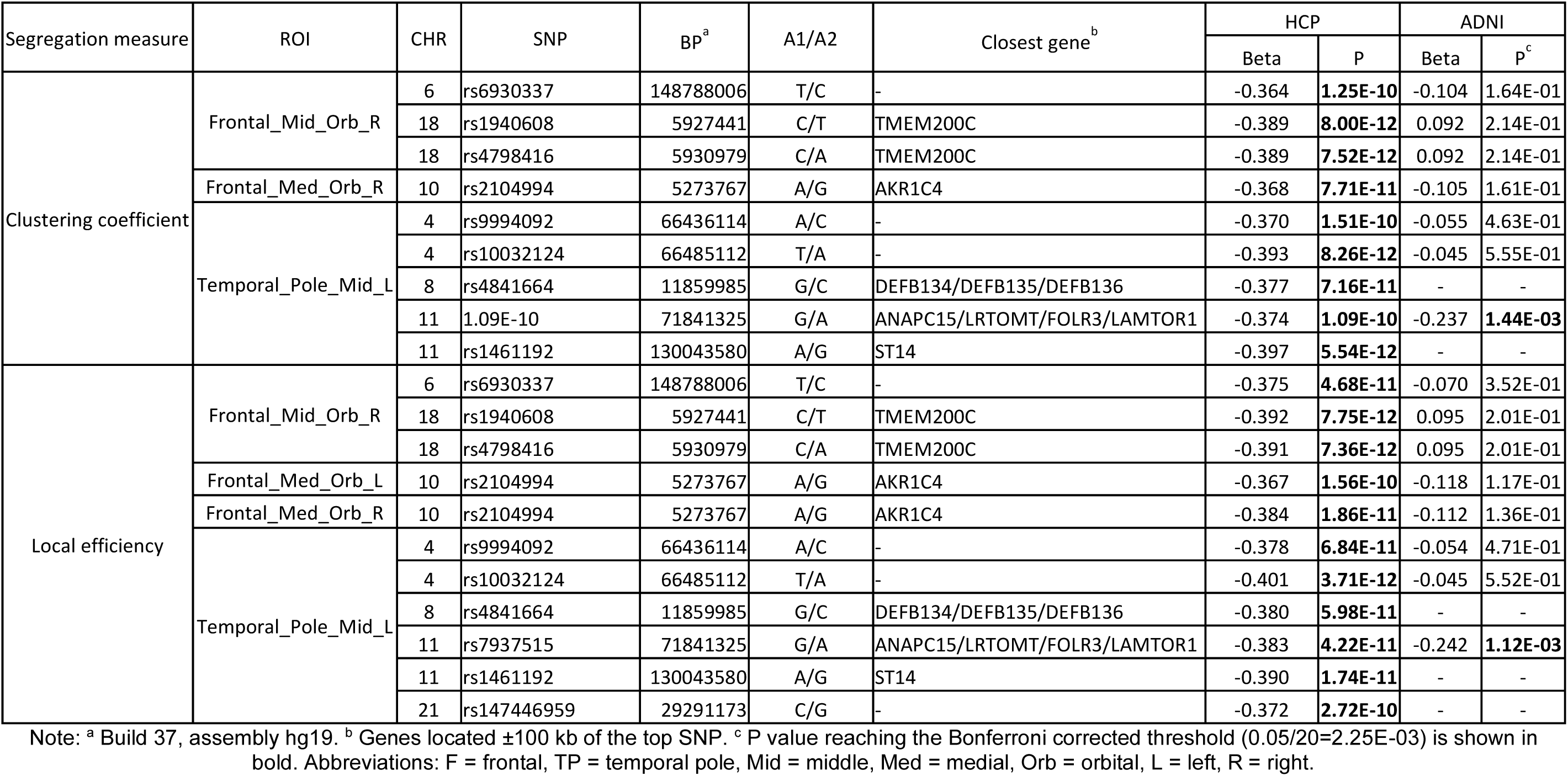
Summary of significant associations between SNPs and segregation measures identified in HCP cohort (Bonferroni corrected p threshold = 2.75E-10) and their replication statistics in ADNI cohort.

### 3.3 Targeted genetic association of segregation in ADNI elderly adults

Given the list of significant findings from the aforementioned GWAS of HCP segregation measures, we further examined their statistical significance in the ADNI cohort to identify brain network relevant genetic variants which were consistent for brain aging. We assessed the associations of 15 out of 20 HCP GWAS findings in ADNI cohort, as three SNPs (rs4841664, rs1461192 and rs147446959 are corresponding to 5 associations in Table 2) were not included in ADNI genotyping data. Associations of rs7937515 with clustering coefficient and local efficiency measured in left middle temporal gyrus were duplicated and validated in ADNI cohort, for this step, we applied the Bonferroni correction of p*<* 0.05/15 = 3.33E-03 (Table 2).

The minor allele G of rs7937515 (rs7937515-G) was associated with lower level of both clustering coefficient and local efficiency, compared to its major allele A (Figure 3). We will discuss the risk effect of rs7937515-G on brain function and dysfunction in the discussion section.

**Figure 3:**
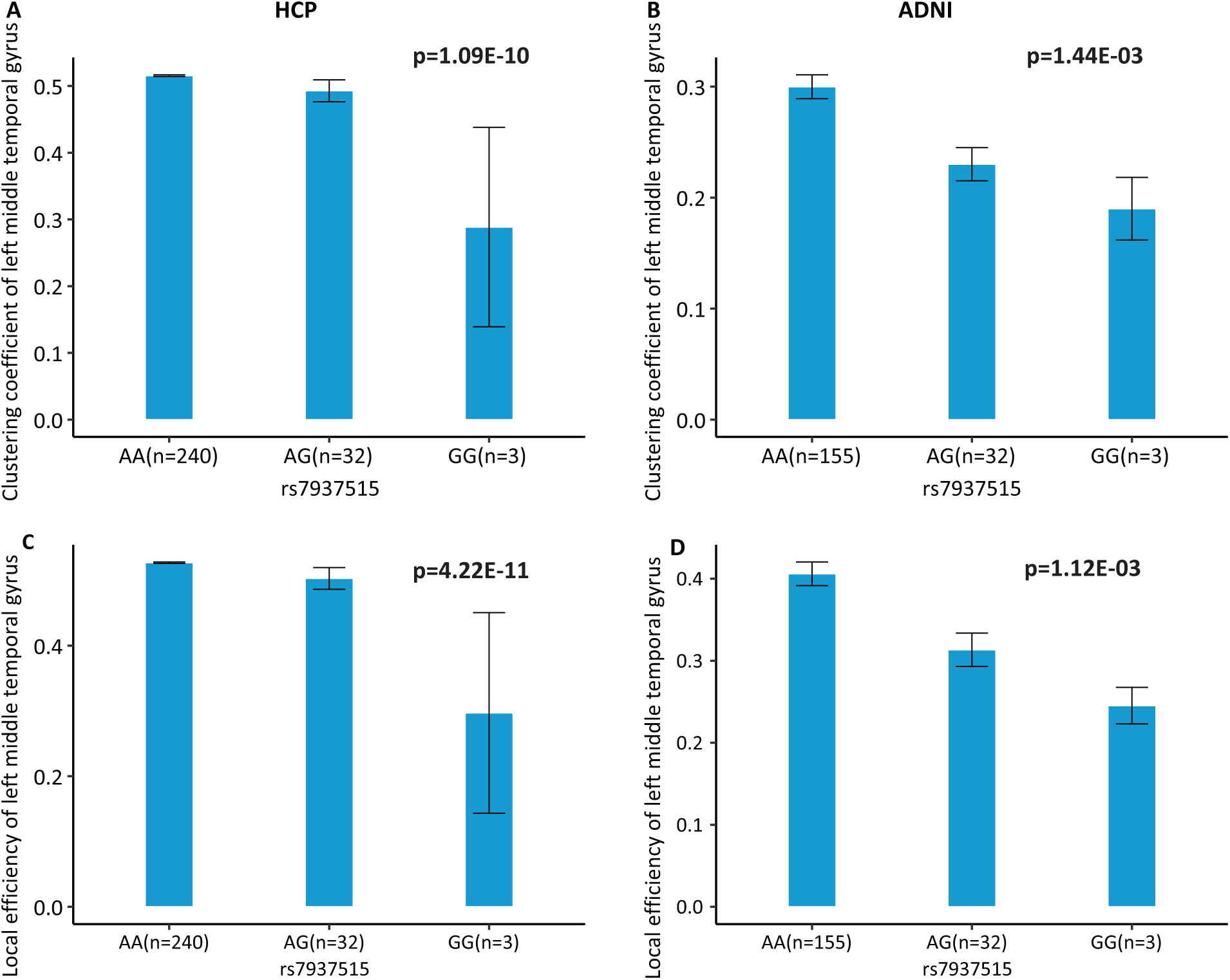
Association of rs7937515 on clustering coefficient and local efficiency of the left middle temporal gyrus. (A-B) Mean clustering coefficient with standard errors are plotted against the rs7937515 genotype groups (i.e., AA, AG and GG) for the HCP and ADNI cohorts, respectively. (C-D) Mean local efficiency with standard errors are plotted against the rs7937515 genotype groups (i.e., AA, AG and GG) for the HCP and ADNI cohorts, respectively. p values are calculated from GWAS (HCP) and targeted analysis (ADNI), and significant p values are marked in bold.

### 3.4 Mediation analysis

According to the genetic association results from the HCP and ADNI subjects, we identified a common genetic finding SNP rs7937515, which was associated with two segregation measures in left middle temporal gyrus (e.g. clustering coefficient and local efficiency). In addition, we extracted two common behavioral and cognitive outcome measures (e.g. BMI and MMSE) by comparing the outcome evaluation methods across the HCP and ADNI databases. Thus, in this section, we had two major focuses: (1) exploring the genetic effect of SNP rs7937515 on outcomes including BMI and MMSE, and gaining deeper insights to the molecular mechanisms of the identified genetic variant, and (2) examining the mediated effect of iQTs (e.g. segregation measures) and illustrating their implicit effects in (1).

To achieve those two goals, mediation analysis of outcome was performed for evaluating both the direct and implicit effects of rs7937515 on outcomes (i.e., BMI and MMSE) through segregation measurements in left middle temporal gyrus. Mediation analysis required the independent variable (i.e., rs7937515) to be significantly associated with both the dependent variable (i.e., BMI or MMSE) and the mediator (i.e., segregation measurements). Below we respectively reported the mediation results analyzed from both HCP and ADNI data.

For the first focus, the minor allele G of rs7937515 was significantly associated with the increased BMI in HCP cohort (p = 1.62E-03; Figure 4A). The same increasing trend was also observed from the ADNI data, although the association between rs7937515 and BMI was not significant (p = 0.22; Figure 4B). For the second focus, Figure 5 showed the result of mediation analysis with BMI as an outcome measure, from which both clustering coefficient and local efficiency of the left middle temporal gyrus demonstrated their intermediate roles between rs7937515 and BMI. There was no significant association between rs7937515 with MMSE in the HCP young adult dataset, so no mediation analysis regarding MMSE was performed. In the ADNI elderly adult dataset, there were no significant associations observed between rs7937515 with BMI nor MMSE; therefore it was not necessary to further examine mediation effects.

**Figure 4:**
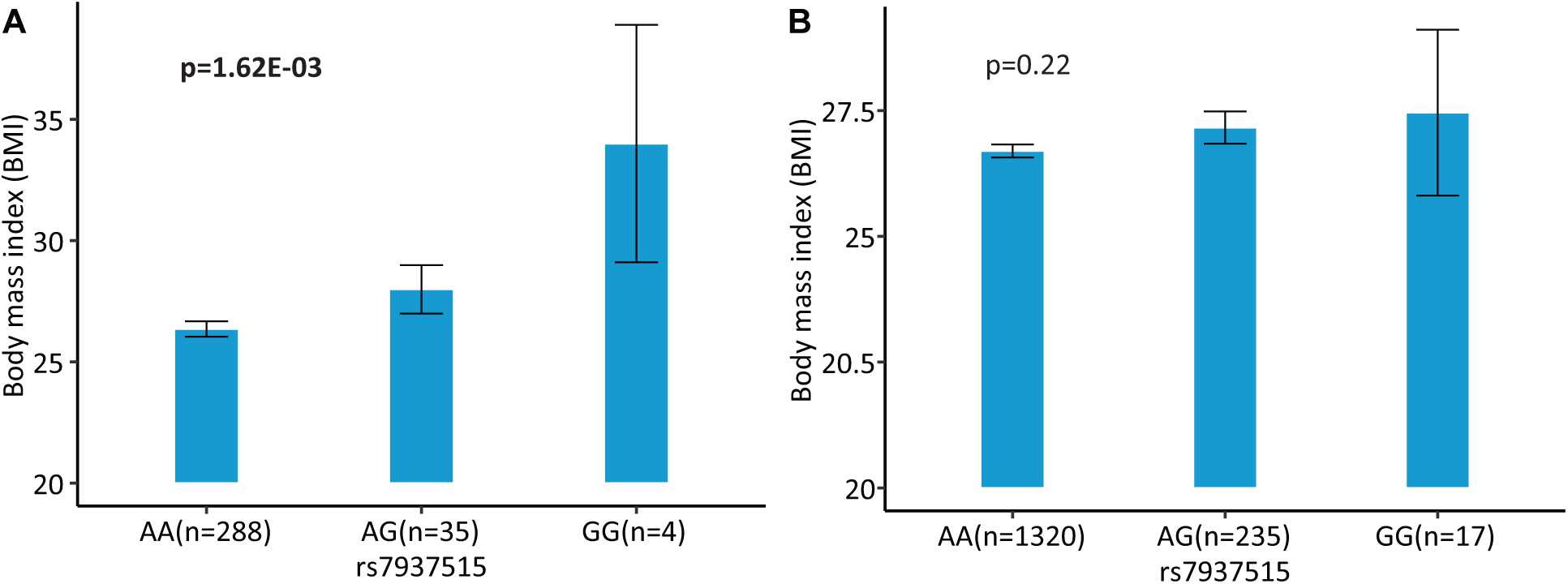
Association of rs7937515 on BMI in the HCP and ADNI cohorts. (A) Mean BMI with standard errors are plotted against the rs7937515 genotype groups (i.e., AA, AG and GG) for the HCP cohort. (B) Mean BMI with standard errors are plotted against the rs7937515 genotype groups (i.e., AA, AG and GG) for the ADNI cohort. p values are calculated from mediation analysis, and significant p values are marked in bold.

**Figure 5:**
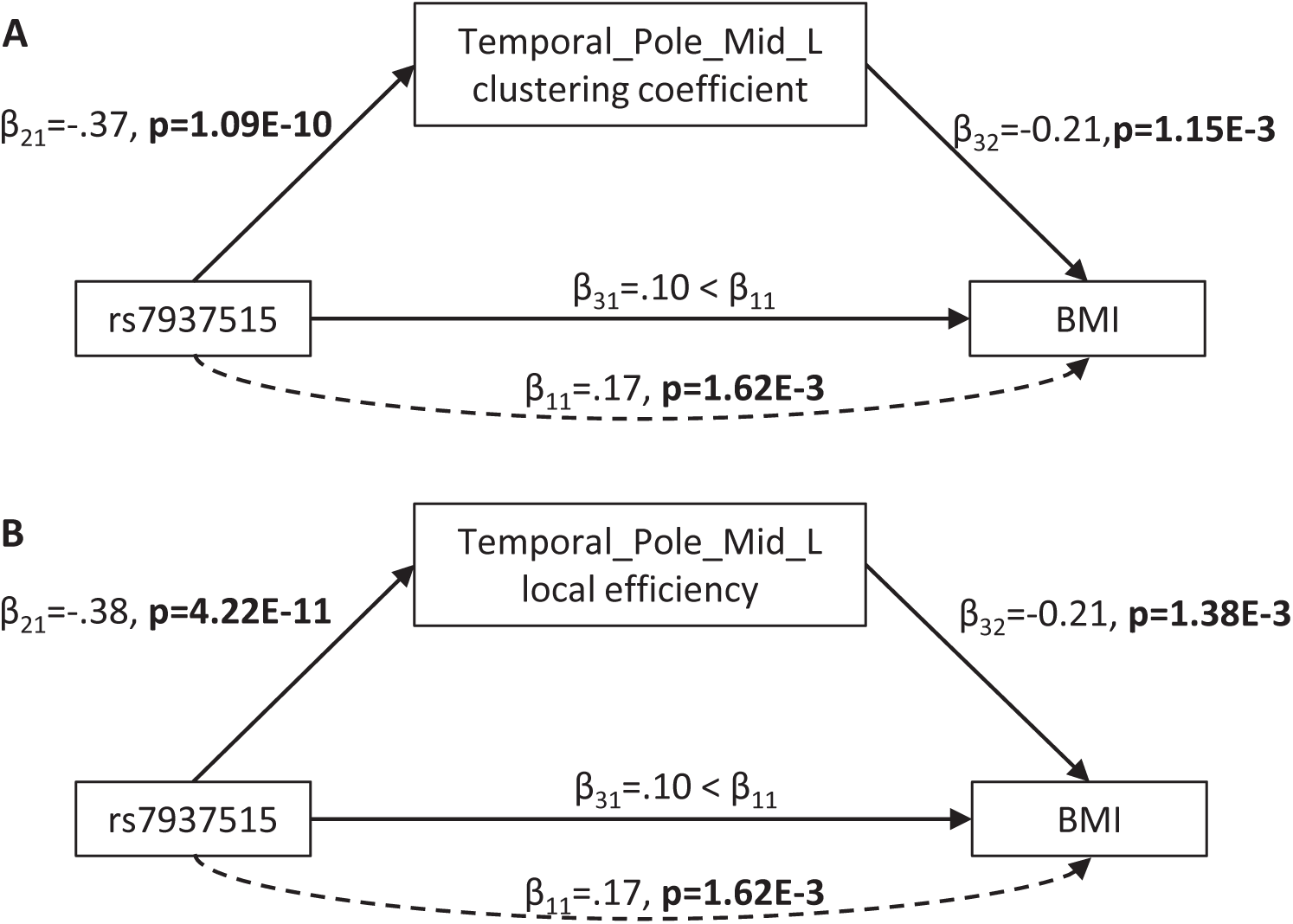
Direct and indirect effect of rs7937515 on BMI through left middle temporal gyrus. (A) and (B) illustrate the statistics of each step from rs7937515 mediation analysis of BMI via left middle temporal clustering coefficient and local efficient, respectively. See Equation 6, 7 and 8 for the definitions of *βs*

### 3.5 Outcome-relevant neuroimaging biomaker discoveries

On one hand, for the HCP cohort, we respectively identified significantly negative associations (p*<* 0.05/4 = 1.25E-02) between BMI with clustering coefficient (p= 3.92E-05) and local efficiency (p= 4.57E-05) measured in left middle temporal gyrus. On the other hand, for the ADNI cohort, we examined the associations between BMI and the above mentioned segregation measures in a pairwise manner, but there was no significant findings satisfying the corrected p threshold. Regarding the relationship between cognitive score (e.g. MMSE) and network segregation measures, there was no significant associations identified for both HCP and ADNI cohorts.

### 3.6 Targeted genetic association of other important network attributes in the left middle temporal gyrus

To review the genetic effect of SNP rs7937515 from different aspects of network connectivity attributes of the left middle temporal gyrus, we assessed the relationship between rs7937515 and additional node-level measures on reported brain ROI (i.e., left middle temporal gyrus) as well as network-level measures in both HCP and ADNI datasets. Table 3 showed association statistics of rs7937515 with segregation, integration, centrality and resilience measures. After correcting for the number of examined network measures (i.e., P*<* 0.05/9 = 5.56e-03), both HCP and ADNI identified significant associations between the targeted SNP with global efficiency (integration) and transitivity (resilience), together with our previous finding that rs7937515 was associated with segregation measures such as clustering coefficient and local efficiency, our results showed the consistent genetic effect of rs7937515 on brain structural network segregation, integration and resilience across aging. Besides the common findings between young and elderly adults, rs7937515 was associated with several other node-level and network-level attributes including network density (integration) and eigenvector centrality (centrality) in HCP data, but not in ADNI. Our results suggested the possible genetic discrepancy for certain brain connectivities in different life stages.

**Table 3:** Associations between rs7937515 and brain network measures.

| Class | QT | Node-level/network-level | HCP |  | ADNI |  |
| --- | --- | --- | --- | --- | --- | --- |
|  |  |  | Beta | P | Beta | P |
| Segregation | Clustering coefficient | Temporal_Pole_Mid_L | -0.37 | **1.09E-10** | -0.24 | **1.44E-03** |
|  | Clustering coefficient | Temporal_Pole_Mid_R | -0.20 | **3.53E-04** | -0.22 | **3.73E-03** |
|  | Local efficiency | Temporal_Pole_Mid_L | -0.38 | **4.22E-11** | -0.24 | **1.12E-03** |
|  | Local efficiency | Temporal_Pole_Mid_R | -0.22 | **7.05E-05** | -0.22 | **2.48E-03** |
|  | Transitivity | Network-level | -0.23 | **3.65E-05** | -0.24 | **9.70E-04** |
|  | Modularity | Network-level | 0.20 | **5.32E-04** | -0.11 | 1.20E-01 |
| Integration | Global efficiency | Network-level | -0.29 | **1.63E-07** | -0.24 | **1.19E-03** |
|  | Density | Network-level | -0.26 | **2.64E-06** | 0.02 | 7.88E-01 |
| Centrality | Betweenness centrality | Temporal_Pole_Mid_L | -0.09 | 1.28E-01 | -0.05 | 5.44E-01 |
|  | Betweenness centrality | Temporal_Pole_Mid_R | -0.06 | 3.24E-01 | -0.03 | 6.62E-01 |
|  | Eigenvector centrality | Temporal_Pole_Mid_L | -0.32 | **9.58E-08** | -0.14 | 6.88E-02 |
|  | Eigenvector centrality | Temporal_Pole_Mid_R | -0.20 | **6.11E-04** | -0.04 | 6.27E-01 |
| Resilience | Assortativity coefficient | Network-level | 0.10 | 1.14E-01 | 0.06 | 4.06E-01 |

Abbreviations: F = frontal, TP = temporal pole, Mid = middle, Med = medial, Orb = orbital, L = left, R = right, QT = quantitative trait.

### 3.7 Hemispheric asymmetry of brain connectome

In this study, we noticed a hemispheric asymmetry of outcome-relevant brain connectivity alterations in the left and right middle temporal gyrus (Table 3). Due to two brain regions (e.g. left and right MTG), two segregation measures (e.g. clustering coefficient and local efficiency) and one outcome measure (e.g. BMI), we applied a Bonferroni corrected p threshold in this section (p*<* 5E-02/8 = 6.25E-03). In the HCP young adult cohort, for the left MTG, we respectively identified significant associations of BMI with clustering coefficient (p= 3.92E-05), and with local efficiency (p= 4.57E-05); for the right MTG, even though there were no significant associations of BMI with clustering coefficient (p= 2.24E-02), and with local efficiency (p= 2.90E-02), both clustering coefficient and local efficiency in left and right MTG showed negative associations with BMI. In the ADNI cohort, as reported in the previous section, network segregation was not associated with BMI, so it was not necessary and proper to conduct analyses regarding ADNI data in this section.

## 4 Discussion

As summarized in Figure 1, prior to GWAS, we first performed heritability analysis for network attributes screening, and only heritable measures of network segregation were treated as iQT for GWAS. Based on experimental outcomes, all of the segregation measures were highly heritable: transitivity and modularity were heritable at network level, clustering coefficient and local efficiency were heritable at all nodes, which suggested segregation measures were suitable for genetic analyses. Then, we performed GWAS of segregation attributes in 275 HCP subjects, and identified several pairwise associations between SNPs and iQTs as listed in Table 2. These GWAS findings were validated in 178 ADNI subjects. As a validation result, we identified several genetic consistency and discrepancy patterns for human connectome in different life stages (as shown in Table 2). As common findings in both HCP young adult and ADNI elderly adult cohorts, rs7937515 was negatively associated with clustering coefficient and local efficiency respectively measured at left middle temporal gyrus. To the best of our knowledge, this was among the first GWAS of human brain high-level network measures across both young and elderly participants. As shown in Figure 3(a,c), the minor allele G of rs7937515 was associated with decreased clustering coefficient and local efficiency of the left middle temporal gyrus in both young and elderly participants. As concluded in (Karwowski et al., 2019; Keown et al., 2017; Rudie et al., 2013; Varangis et al., 2019), the weakness of segregated network connectivity (e.g. modularity, clustering coefficient, and local efficiency) was linked to multiple brain disorders such as age-related cognitive declines and autism spectrum disorder. Thus, our GWAS findings for HCP young adults demonstrated that rs7937515 played a risk effect on human network segregation. This neurorisk effect was also confirmed in a targeted genetic association analysis for ADNI elderly participants (as shown in Figure 3 (b,d)), these common discoveries between HCP and ADNI datasets suggested a consistent genetic risk effect across young and old life stages.

This study was further conducted by performing several post-hoc analyses in the following three aspects (shown as bottom sections in Figure 1): 1) examining genetic mechanisms for common outcome measures in the HCP and ADNI studies, and elucidating the mediated effect of iQTs for such outcome-relevant genetic association, 2) discovering outcome-relevant imaging biomarkers, and 3) exploring the genetic mechanisms of other important complex-network attributes.

For the first aspect, our goal was to elucidate the neurobiological pathway from SNPs to brain connectome, and to phenotypic outcome. In addition, we aimed to discover the role of iQTs in the outcome-relevant genetic associations by performing mediation analyses in both HCP and ADNI datasets. For the HCP young participants, we identified that BMI was positively associated with rs7937515 in the first step of mediation analysis, demonstrating a risk effect. rs7937515 located in the regions of *FAM86C1*/FOLR3 was previously discussed in literatures (Gao, 2017; Gao et al., 2015) and positively linked to waist circumference in the meta-analysis based on the Insulin Resistance Atherosclerosis Family Study (IRASFS) (Palmer et al., 2015), which was designed to investigate the genetic and environmental basis of insulin resistance and adiposity. *FAM86C1* (Family With Sequence Similarity 86 Member C1) and *FOLR3* (Folate Receptor Gamma) had been reported for their associations with various weight-related phenotypes such as bone mineral density (Li et al., 2019) and BMI (Hair, 2014; Mrozikiewicz et al., 2019), which closely related to osteoporosis (Li et al., 2019; Mrozikiewicz et al., 2019) and obesity (Gómez-Ambrosi et al., 2004). In the second and third steps of mediation analysis, we illustrated that BMI was indirectly influenced by rs7937515 (Figure 4 and 5), and iQTs such as clustering coefficient and local efficiency measured at the left middle temporal gyrus respectively played a mediating role. We also examined the genetic association with MMSE, but no evidence indicated any genetic associations to MMSE. In contrary, for the ADNI elderly participants, neither significant associations between rs7937515 and BMI nor MMSE were identified in the first step of mediation analysis, so there was not a necessary to examine mediated effect in this dataset. Our results demonstrated a disappearance of outcome-relevant genetic effect in the elderly participants, this discrepancy from young to elderly participants might due to the dominated influences from life style, environment or other non-genetic factors.

For the second aspect, our goal was to reveal the mapping between connectivity alterations and phenotypic outcome, and discover outcome-relevant imaging biomarkers. For young adult participants, segregation measures (e.g. clustering coefficient or local efficiency measured at left middle temporal gyrus) previously demonstrated their potential to play a mediating role in genetic association discoveries, in this step, we focused on examining their direct associations to the outcomes. Thus, we performed a targeted association analysis between the mentioned segregation measures and the common outcomes (e.g. BMI or MMSE) evaluated in both HCP and ADNI studies (Table 2) by employing linear regression models. For the young participants, clustering coefficient and local efficiency measured at left middle temporal gyrus were negatively associated with BMI. Similar observation was obtained in (Chen et al., 2018) which linked lower structural network segregation to obesity (higher BMI). Our findings suggested that there was a mapping between brain network segregation attributes and human physical conditions, and segregation features of the left middle temporal gyrus showed their potential as neuroimaging biomarkers to detect human development and BMI related health issues. For elderly adult participants, no significant associations were identified between segregation measures and any outcomes, which suggested an interesting topic for further explorations.

Multiple regression analyses demonstrated that middle temporal gyrus was linked to weight-related issues. For example, Veit et al. (Veit et al., 2014) and Gómez-Apo et al. (Gómez-Apo et al., 2018) revealed that BMI, visceral fat and age were negatively associated with cortical thickness of the left middle temporal gyrus, Ou et al. (Ou et al., 2015) indicated that greater adiposity was associated with lower gray matter (GM) volumes in the middle temporal gyrus, Yokum et al. (Yokum et al., 2012) found positive correlation between BMI and white matter (WM) volume in the middle temporal gyrus, Rapuano et al. (Rapuano et al., 2016) illustrated left middle temporal gyrus was detected significantly greater activation in response to food commercials than to non-food commercials, Salzwedel et al. (Salzwedel et al., 2019) concluded that maternal adiposity influenced neonatal brain functional connectivity in middle temporal gyrus, and Peven et al. (Peven et al., 2019) identified that cardiorespiratory fitness was negatively associated with functional connectivity in the right middle temporal gyrus. To the best of our knowledge, our investigations for the association between structural connectivity in the middle temporal gyrus and BMI was among the first weight-related studies with high-level imaging features measured from structural network connectivity, and our results confirmed several previous findings analyzed from thickness data, T1-weighted MRI data, and fMRI data.

For the third aspect, since there was an emerging interest in understanding the segregation and the integration of brain networks (Cohen and D’Esposito, 2016; Mohr et al., 2016) as well as other important network attributes such as centrality (Zuo et al., 2012) and resilience (Karwowski et al., 2019), our goal was to expand our focus on comprehensively discussed segregation attributes to a more complete set of network attributes including segregation, integration, centrality and resilience. For both node level network attributes measured at left and right middle temporal gyrus and global network attributes, we applied targeted genetic association analyses on global efficiency and density (integration, network level), betweeness and eigenvector centrality (centrality, node level) and assortativity coefficient (resilience, network level) of the structural connectivity. We identified several pairwise associations between rs7937515 and these network attributes in both HCP and ADNI datasets (Table 3), and noticed a significant association between rs7937515 and global efficiency in both datasets, which suggested that rs7937515 was involved into the dynamic fluctuations of segregation and integration of neural information. This finding partially answered an elusive question of revealing genetic basis for brain mechanisms of balancing network segregation and integration. Another worth noting finding was that rs7937515 was associationed density and eigenvector centrality respectively in our targeted analyses, while such associations were vanished in elderly participants, which suggested inconsistent genetic influences across different life stages.

With the awareness of the hemispheric asymmetry of network organization, a genetic basis to explain the differences in connectome between two hemispheres were under discovered. In this work, we identified an obvious inconsistency of genetic influences on human connectome in different brain hemispheres (Table 3). As reported in several recent studies (Jiang et al., 2019; Shu et al., 2015; Tian et al., 2011), the topological organizations of structural networks were not uniformly affected across brain hemispheres, which lead to a non-uniformly distributed destruction on neural network of the left and right hemispheres. Our finding gave an explanation from the point-view of genetics, with the potential for further investigations as many of the destruction on neural network (as iQT) were linked to cognitive and behavioral functions and dysfunctions, and their genetic mechanisms were still under discovered.

## 5 Conclusions

In this work, we constructed the structural network connectivity, extracted complex-network attributes and examined the heritability of network segregation measures. Then, we revealed a novel association between the minor allele (G) of rs7937515 and decreased network segregation measures of the left middle temporal gyrus across HCP young participants and ADNI elderly participants, which demonstrated a consistent genetic risk effect on brain network connectivity across lifespan. We elucidated the neurobiological pathway from SNP rs7937515 and genes *FAM86C1*/*FOLR3* to brain network segregation, and to BMI. In such pathway, we concluded a genetic risk effect on BMI due to their positive association, examined the mediated effect of network segregation measures, and discovered network segregation of the left middle temporal gyrus as BMI-related neuroimaging biomarkers by identifying a negative association between them. We also examined genetic associations of a more complete set of important network attributes including integration, centrality and resilience, and demonstrated the consistency and discrepancy in genetic associations in brain aging. At last, we illustrated hemispheric asymmetry of network organization in both datasets in the aspect of genetic effect.

In sum, this study performed a systematic analysis that integrated genetics, connectomics and phenotypic outcome data, with the goal of revealing biological mechanisms from the genetic determinant to intermediate brain connectomic traits and to the BMI phenotype at two different life stages. We identified the genetic effect of rs7937515 on human brain network segregation in both young and elderly participants and on BMI in young adult cohort. Our findings confirmed several previous genetic and imaging biomarker discoveries. Moreover, we provided outcome-relevant genetic insights in the aspect of complex-network attributes of human brain connectome. Similar analytical strategies can be applied to future integrative studies linking genomics with connectomics, including the analyses of structural and functional connectivity measures, additional network attributes, longitudinal or dynamic connectomic features, as well as other types of brain imaging genomic data.

## Ethics Approval

This research is conducted under the regulation of Institutional Review Boards (IRB) and the research subject informed consent process at University of Pennsylvania, USA. Study subjects gave written informed consent at the time of enrollment for data collection and completed questionnaires approved by each participating site’s IRB. The authors state that they have obtained approval from the Alzheimer’s Disease Neuroimaging Initiative (ADNI) and Human Connectome Project (HCP) Data Sharing and Publications Committee for use of the data.

## Consent to participate

Informed consent was obtained from all individual participants included in the study.

## ADNI Acknowledgements

Data collection and sharing for this project was funded by the Alzheimer’s Disease Neuroimaging Initiative (ADNI) (National Institutes of Health Grant U01 AG024904) and DOD ADNI (Department of Defense award number W81XWH-12-2-0012). ADNI is funded by the National Institute on Aging, the National Institute of Biomedical Imaging and Bioengineering, and through generous contributions from the following: AbbVie, Alzheimer’s Association; Alzheimer’s Drug Discovery Foundation; Araclon Biotech; BioClinica, Inc.; Biogen; Bristol-Myers Squibb Company; CereSpir, Inc.; Cogstate; Eisai Inc.; Elan Pharmaceuticals, Inc.; Eli Lilly and Company; EuroImmun; F. Hoffmann-La Roche Ltd and its affiliated company Genentech, Inc.; Fujirebio; GE Healthcare; IXICO Ltd.; Janssen Alzheimer Immunotherapy Research & Development, LLC.; Johnson & Johnson Pharmaceutical Research & Development LLC.; Lumosity; Lundbeck; Merck & Co., Inc.; Meso Scale Diagnostics, LLC.; NeuroRx Research; Neurotrack Technologies; Novartis Pharmaceuticals Corporation; Pfizer Inc.; Piramal Imaging; Servier; Takeda Pharmaceutical Company; and Transition Therapeutics. The Canadian Institutes of Health Research is providing funds to support ADNI clinical sites in Canada. Private sector contributions are facilitated by the Foundation for the National Institutes of Health (www.fnih.org). The grantee organization is the Northern California Institute for Research and Education, and the study is coordinated by the Alzheimer’s Therapeutic Research Institute at the University of Southern California. ADNI data are disseminated by the Laboratory for Neuro Imaging at the University of Southern California.

## HCP Acknowledgements

Data were provided by the Human Connectome Project, WU-Minn Consortium (Principal Investigators: David Van Essen and Kamil Ugurbil; 1U54MH09-1657) funded by the 16 NIH Institutes and Centers that support the NIH Blueprint for Neuroscience Research; and by the McDonnell Center for Systems Neuroscience at Washington University.

## Authorship Confirmation Statement (CRediT Format)

**Shan Cong:** Conceptualization, Methodology, Software, Formal analysis, Validation, Writing - Original Draft. **Xiaohui Yao:** Formal analysis, Validation, Writing - Review & Editing. **Jingwen Yan:** Data Curation, Resources Writing - Review & Editing. **Linhui Xie:** Data Curation, Software, Writing - Review & Editing. **Li Shen:** Supervision, Conceptualization, Methodology, Writing - Review & Editing.

---

**Supplementary Table:**
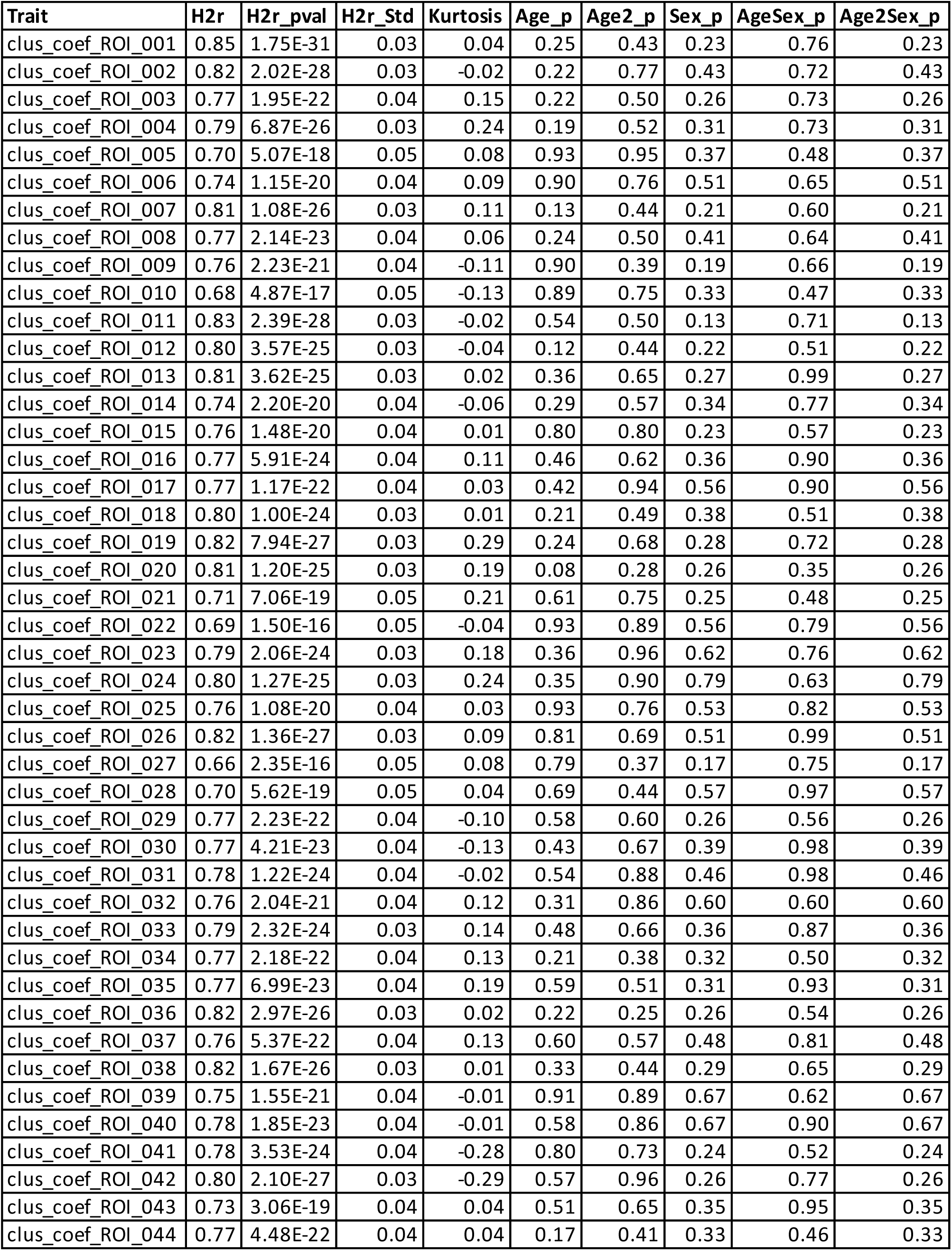
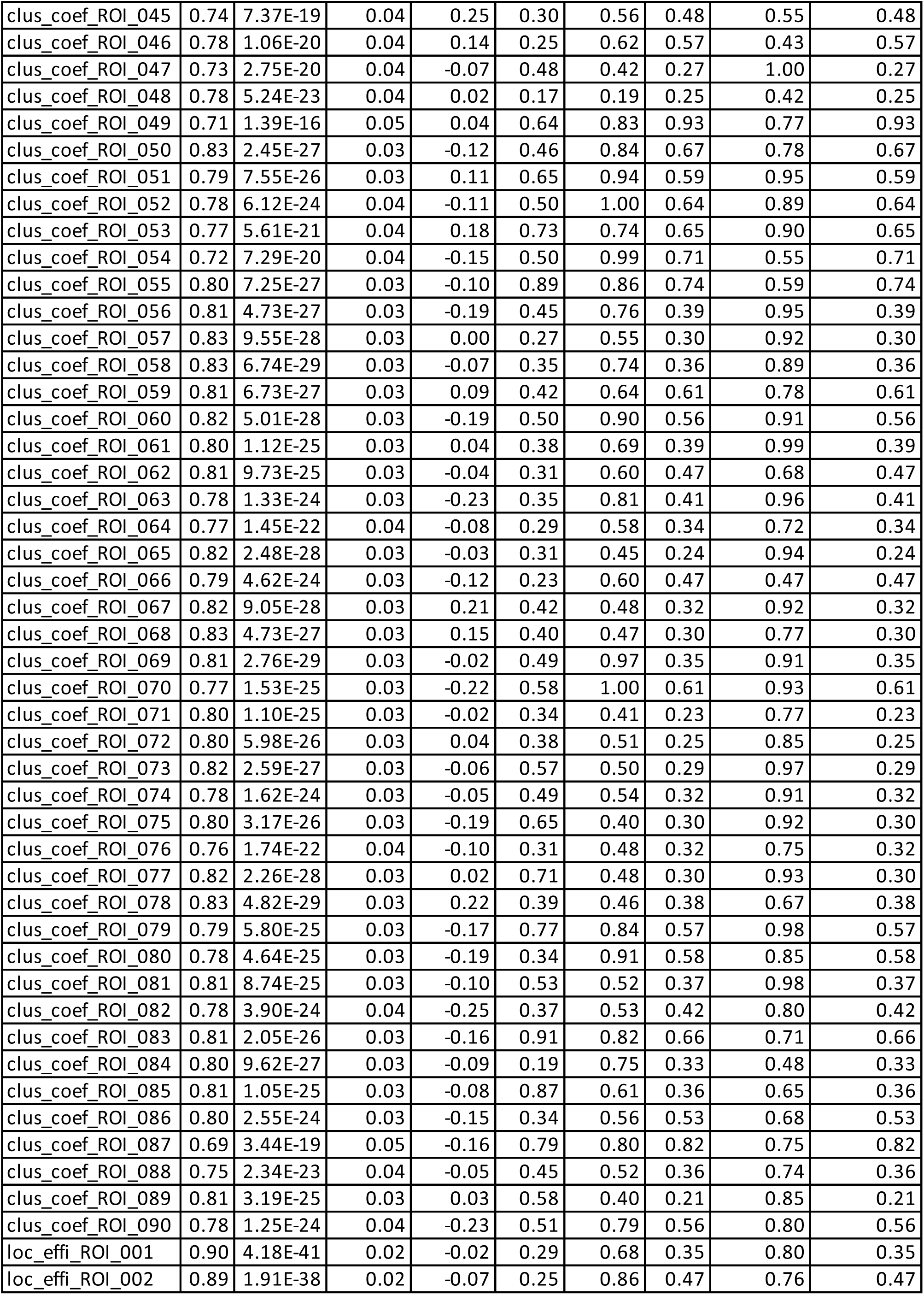
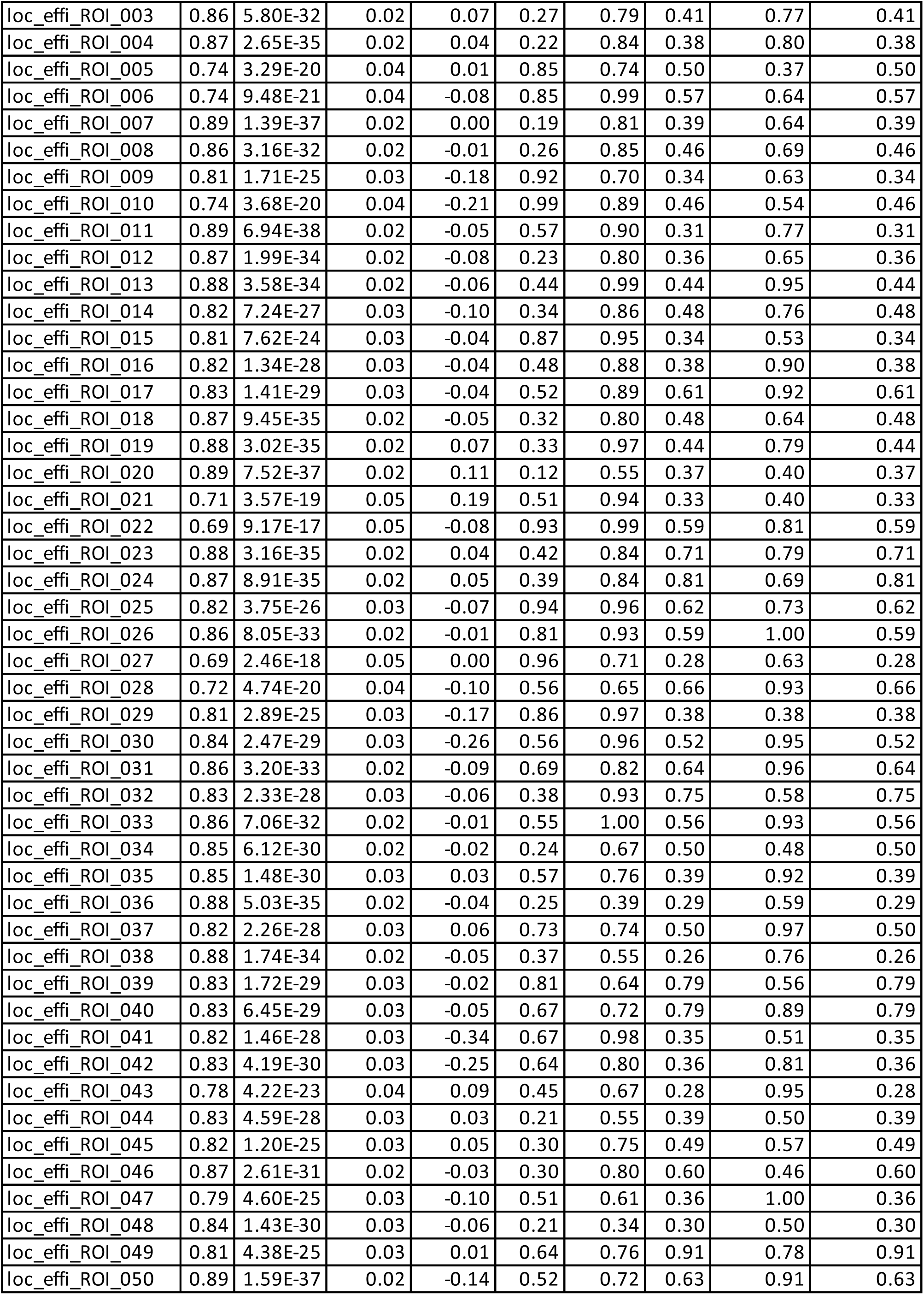
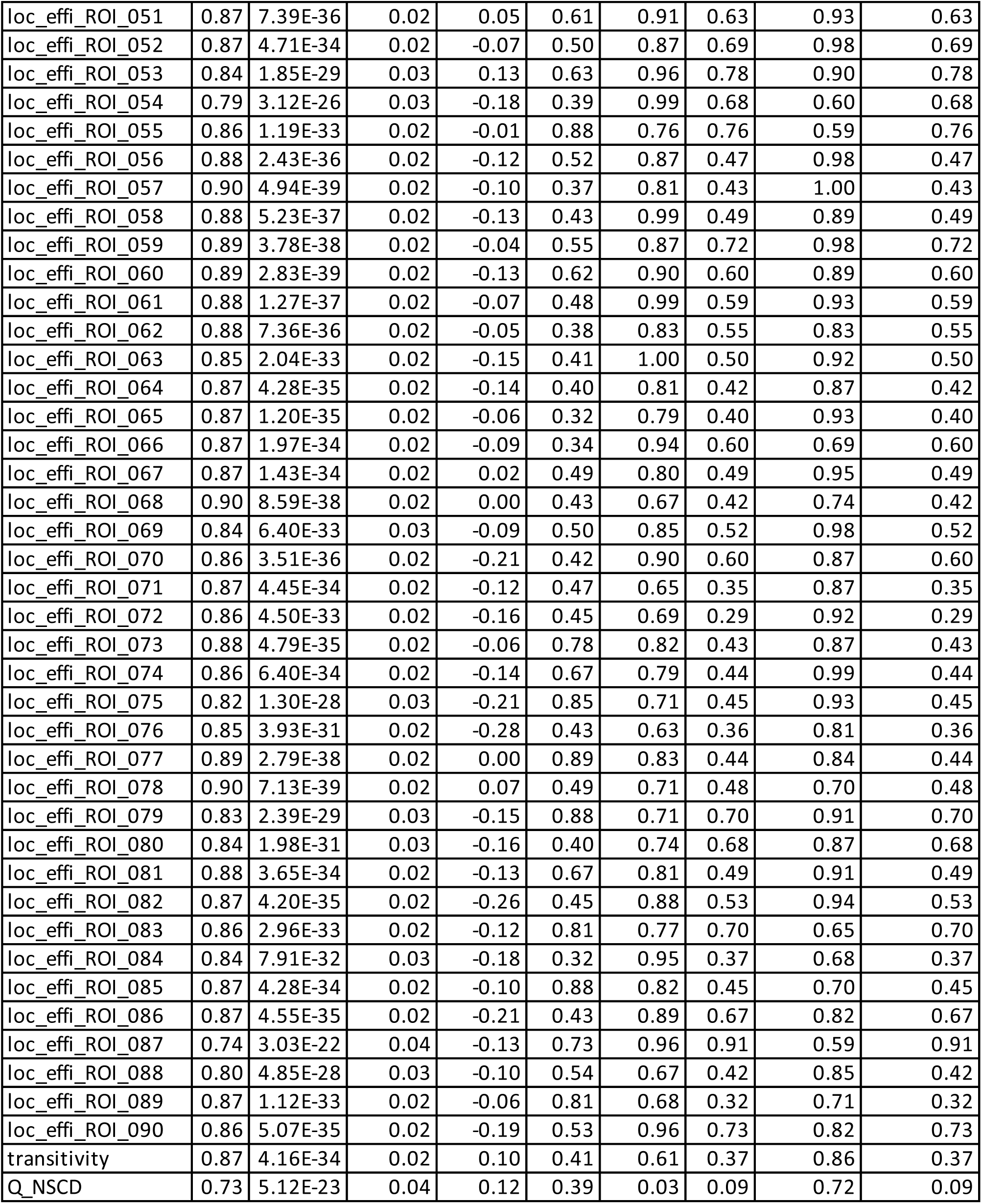
Results of heritability analysis.

